# Distributed Collaboration for Data, Analysis Pipelines, and Results in Single-Cell Omics

**DOI:** 10.1101/2024.07.30.605714

**Authors:** Alexandre Hutton, Lizhuo Ai, Jesse G. Meyer

## Abstract

Single-cell omics data analysis pipelines are complicated to design and difficult to share or reproduce. We describe a web platform that enables no-code analysis pipeline design, simple computing via the Open Science Grid, and sharing of entire data analysis pipelines, their input data, and interactive results. We expect this platform to increase the accessibility and reproducibility of single-cell omics.

## Main

Analyses for single-cell omics data are developing quickly; ensuring that analyses produce consistent and reliable results is important for the field to build confidence in published results. Given that analytical pipelines can have dozens of parameters, communicating these values via traditional publication venues is cumbersome and makes it difficult for others to implement an identical analysis. Analyses are further biased by the preference of the group implementing them^1^. Even when analytical code is included with publications, results can be affected by differences in software dependencies and computing environments^2^. Addressing these issues requires technical knowledge that may not be present in every research group or might be at odds with research priorities.

Rather than expecting each group to obtain software development skills or changing current incentive structures, we instead propose to lower the threshold for making analyses reproducible and transparent to users without impeding research progress. We have developed the Platform for Single-Cell Science (PSCS, https://pscs.xods.org) to help researchers produce research products that bundle data, analysis, results, and communication together. There exist other sites (e.g., ezSC^3^, ICARUS^4^, Cellar^5^) that offer part of the solution by providing fixed pipelines, which are helpful for running a specific analysis (or set of analyses). However, analytical methods change with time and fixed pipelines are difficult to maintain over long periods of time. More importantly, different experiments can require different treatments of their data (e.g., differing quality-control parameters) and a one-size-fits-all analysis is unlikely to be sufficient. Ambitious platforms such as Galaxy^6^ are fantastic resources but lack important features to promote analytical development. The web platform PSCS enables a flexible no-code data analysis and allows for the distribution of data, pipelines, and results in parallel with existing publishing venues.

PSCS includes several features that are designed to increase the accessibility of single-cell omics data analyses, manage projects, and make results publicly viewable. A PSCS project encapsulates data, analyses, and results **(Supplemental Video 1)**. Researchers create a project, upload data, and invite collaborators to join. Analytical pipelines are developed within a project using the no-code pipeline designer, which currently offers the functionality of Scanpy^7,8^, MiloPy^9,10^, and CAPITAL^11^**(Supplemental Video 2)**. Individual functions are available as nodes, each node can have its parameters modified, and nodes from different modules can be used in any order assuming prerequisites are met. To help guide users, PSCS can perform validation before running to verify that node requirements are met. When run, analysis inputs are matched with user-selected data, sent to the Open Science Grid^12-15^ for computing, and executed through a Singularity^16^ container to ensure the same results are obtained on any cluster. The same analysis can be run on different datasets simply by selecting them, allowing for pipeline reuse without having to modify the analysis.

Results from executed pipelines are brought back to PSCS and made available for viewing by the research group. Once researchers are satisfied with a project, it can be published in one of two ways: fully public or secured through a password to allow anonymous viewing during peer review **(Supplemental Video 3)**. The latter allows reviewers to audit a project while keeping it private to prevent the work from being scooped. **Figure 1A** shows the typical workflow from a project’s creation to its publication, and eventual re-use in new projects. The structure encourages the iterative nature of experimentation, and by simplifying the re-use of both data and previously developed analyses we can iterate across independent research groups more easily. The platform is implemented with current technologies (**Figure 1B**). At no point does the user need to worry about how to distribute their data, whether their analysis is version-controlled, or whether their results are reproducible. Data is made available for download upon publication, analyses are kept constant after being saved, and reproducibility is handled by the server. Additionally, pipeline development is aided by a validator to detect problems with the pipeline and make suggestions to fix them (**Figure 1C**). Parts of the pipeline designer are outlined in **Supplemental Figure 1**.

**Figure 1.**
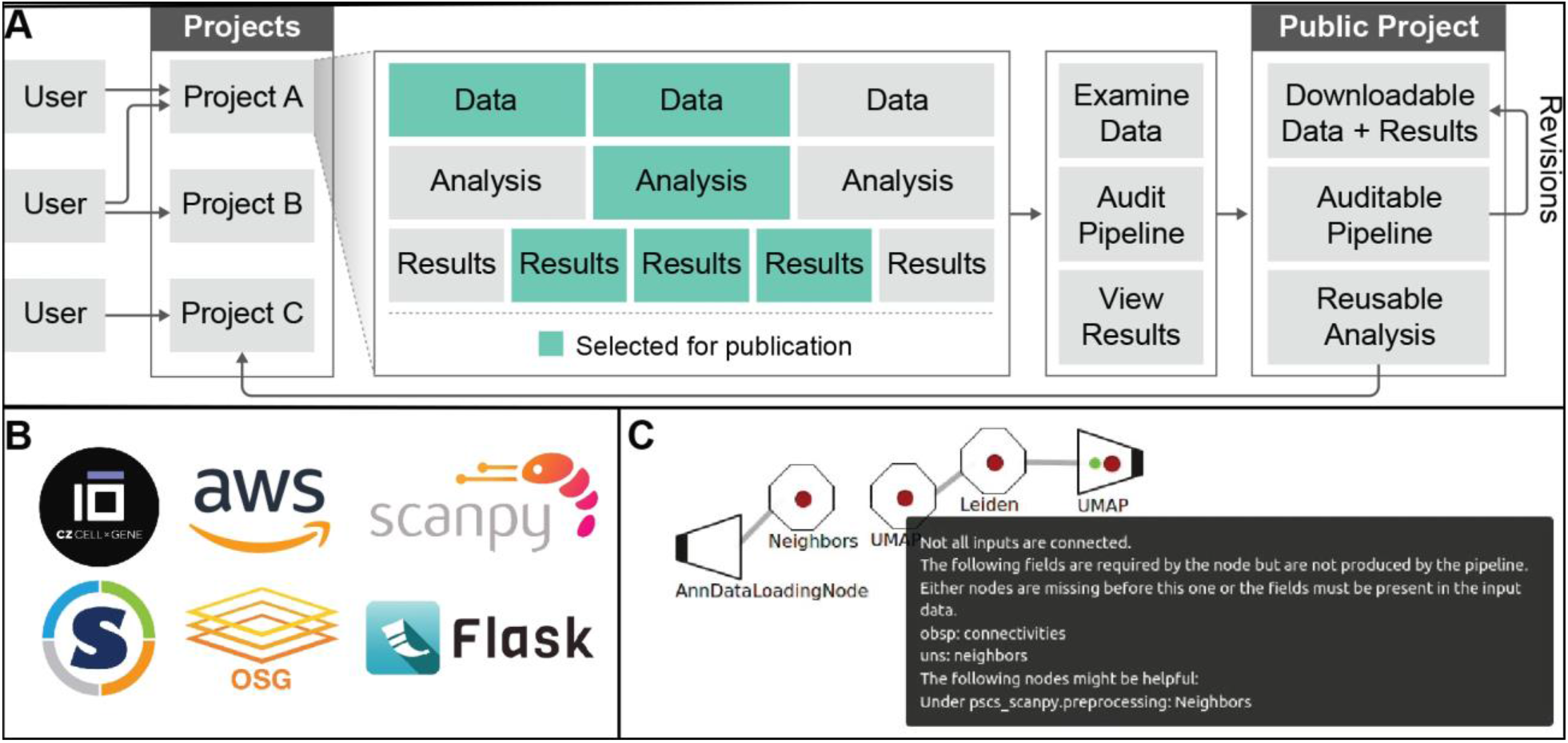
Overview of PSCS. (**A**) Workflow for the development of a PSCS project. Once published, data/analyses are readily reusable by other groups. (**B**) Software packages and technologies used in PSCS. (**C**) Example of validator suggestions during pipeline development.

To exemplify the basic functionality of PSCS including sharing data, analytical pipeline, and results, we recreate the results from the Scanpy PBMC tutorial^17^(**Supplementary Figure 2**, https://pscs.xods.org/p/aE6PG). This pipeline demonstrates all common qualitative steps in single-cell data analysis, including normalization, scaling, dimension reduction, clustering, visualization and marker gene extraction. The project also highlights interactive results exploration, achieved by integrating the CellxGene app^18^.

**Figure 2.**
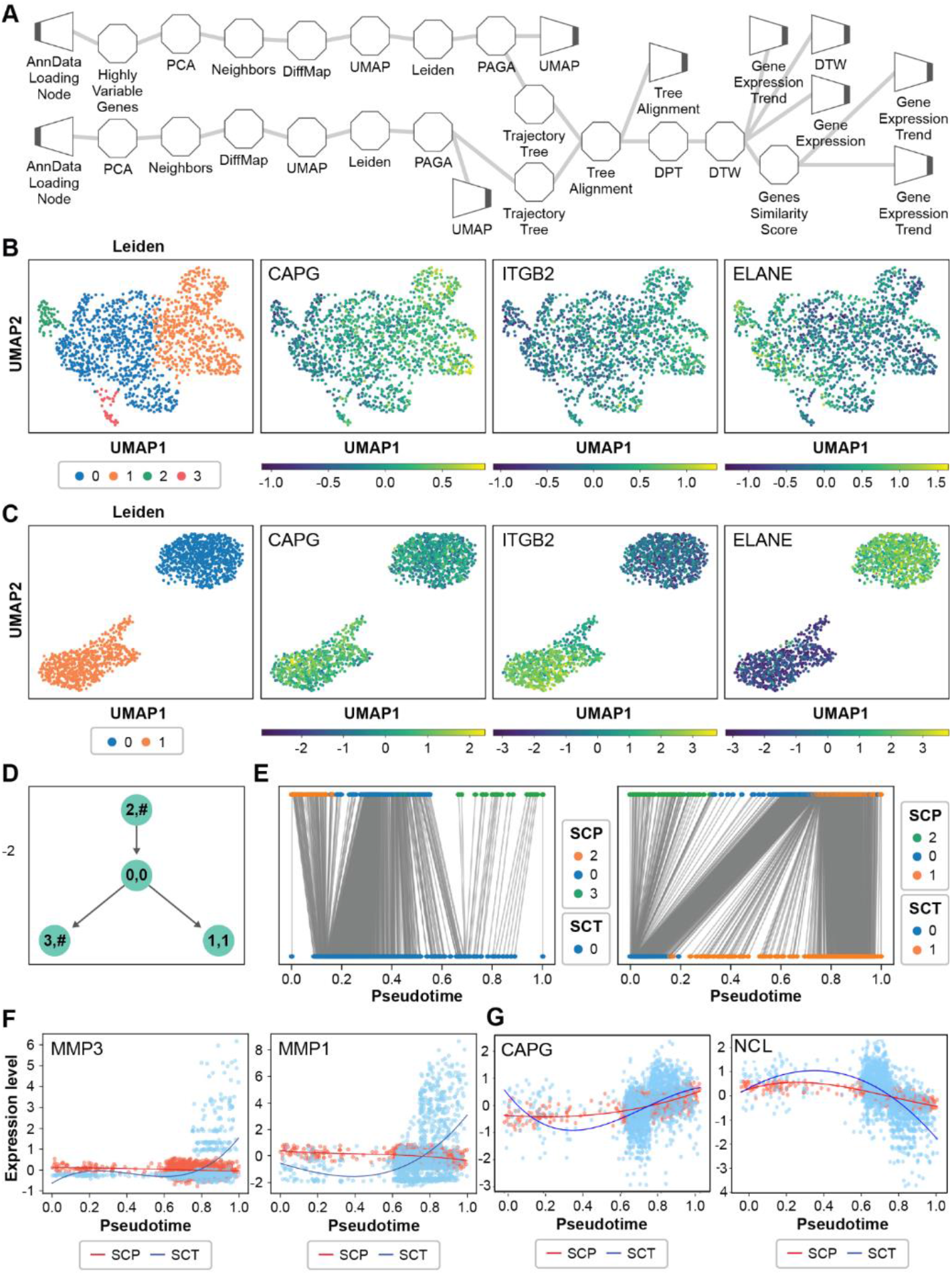
CAPITAL workflow applied to data provided with the SCoPE2 project. **(A)** Schematic of the CAPITAL pipeline for aligning proteomic (top) and transcriptomic (bottom) datasets. Looking from left to right, the first trapezoidal nodes represent data input to the pipeline, the octagonal nodes represent processing nodes, and the trapezoidal nodes on the far right represent pipeline outputs that can be figures or data objects. Data is loaded into the separate paths, preprocessed separately, and combined in the TreeAlignment node. **(B)** UMAP of proteomic data with cluster labels produced by the Leiden algorithm, and quantities for selected proteins. **(C)** As **(B)**, but for the transcriptomic data. **(D)** Alignment tree generated by CAPITAL. The left path or right path are two possible alignments that either include cluster 1 or 3 from the proteomics data. **(E)** Pseudotime alignments for the datasets, one alignment each for the branches of the tree in panel **(D). (F)** Pseudotime trajectories for low-agreement genes. **(G)** Pseudotime trajectories for two selected agreeing genes, NCL and CAPG.

To demonstrate the multi-omic data integration utility of PSCS, we have put together a currently public project^19^ that is described in **Figure 2**. The project uses data from Specht *et al*.^19^ but combines PSCS nodes from the Scanpy and CAPITAL packages to produce novel insights through multi-omic data integration. **Figure 2A** shows the pipeline, with separate paths for the proteomic and transcriptomic data, highlighting the flexibility of PSCS pipelines. Using this analysis, we were able to reproduce the paper’s finding that there are two groups of cell types (monocytes or macrophages). **Figure 2B** and **2C** show the distribution of quantities for proteomics (B) and transcriptomics (C). Trajectory inference and pseudotime alignment with dynamic time warping were performed with CAPITAL (**Figures 2 D-G**), and demonstrates which transcripts and proteins change differently (**Figure 2F**) or the same (**Figure 2G**) as cells transition to macrophages. For example, both the protein and RNA level for CAPG appear to increase across aligned pseudotime. We can see an overview of the project on its publication page (https://pscs.xods.org/p/SzFKQ), including the data, analytical results, and auditable pipeline. PSCS users that would like to use the pipeline on their own work can import it directly to their projects. Both data and results can be directly downloaded for offline examination.

Besides facilitating the use of existing packages and re-use of published pipelines, PSCS aims to encourage researchers to develop new analytical packages and deploy them. We have built a node template^21^ that researchers can use to convert their work into a format that can be fully integrated into PSCS. The PSCS node template represents individual pipeline nodes and is intended to represent a “unit” of analysis. At its core, the template only requires developers to define input parameters and operations on data from the input data object. Managing user parameter settings, connecting to other nodes, and returning files to the user are all handled by PSCS. The intent is to give the field the tools to converge on both the best analytical practices and on robust results by removing barriers that slow down development and distribution. Researchers that would benefit from quickly using established analyses can do so, while those that develop the analyses benefit from the distribution of analytical packages to a broader audience.

In summary, single-cell omics research is a rapidly developing field, but ensuring reliable and reproducible analyses is difficult due to complex code-based pipelines and variations in computing environments. To address this, we introduced PSCS, a web platform that simplifies single-cell omics data analysis while maintaining high flexibility. PSCS offers a no-code interface for designing analysis pipelines using pre-built modules from popular packages like Scanpy.

Users can drag-and-drop these modules to build custom pipelines without writing code. Analyses are then executed on the Open Science Grid using containers, ensuring consistent results. The platform’s functionalities are showcased by reproducing published results and integrating data from multiple sources (proteomic and transcriptomic) using a custom PSCS pipeline. Furthermore, PSCS empowers researchers to develop and share their own analytical tools, fostering collaboration and the development of robust analysis methods within the field. Overall, PSCS offers a user-friendly and comprehensive solution for single-cell omics data analysis, and we expect the platform to facilitate the production of reliable results and accelerate the promotion of seamless collaborative work across multiple research groups.

## Acknowledgements

We thank Dasom Hwang for graphic design support. This work was supported by the NIH NIGMS (R35 GM142502). This research was enabled using services provided by the OSG Consortium^12-15^, which is supported by the National Science Foundation awards #2030508 and #1836650. LA is supported by the California Institute for Regenerative Medicine (CIRM) Scholar program (CIRM EDUC4-12751).

## Author Contributions

Conceptualization, AH and JGM; Methodology, AH and JGM; Software, AH; Validation, AH, LA; Formal Analysis, AH and JGM; Investigation, AH and JGM; Resources, AH and JGM; Data Curation, AH; Writing - Original Draft, AH and JGM; Writing - Reviewing and Editing, AH, LA, JGM; Visualization, AH and JGM; Supervision, JGM; Project Administration, AH and JGM; Funding Acquisition, JGM

## Ethics Declarations

Competing interests

The authors declare no competing interests.

## Online Methods

### Site Implementation

The site has a standard implementation, using the Flask^22^ web framework for processing web requests. Web pages are rendered by Flask using the Jinja2^23^ template engine. Interactive portions of the site are implemented in JavaScript, with the exception of CellXGene Annotate^18^ which is containerized via Docker, wrapped in a Flask application, and run as a microservice. The database is managed with SQLite.

### Pipeline Node Template

Pipelines are constructed from individual nodes. Nodes come in three basic types: input, output, and intermediate. Inputs nodes are matched with files to allow them to import data into the pipeline. Output nodes are used to identify the output files that should be registered with PSCS and made available to the user. Intermediate nodes don’t interact with project files, but data in the pipeline, apply a set of operations, and make the result available to downstream nodes. The generic node template has several sections. Class variables are used to identify important parameters, node requirements, and node effects. Parameters that are labeled as important are displayed when the user modifies the parameters; other parameters are accessed by expanding the parameter modification window. Node requirements specify what variables should be available in the node’s input Annotated Data object in order for the node to run correctly. For example, Scanpy’s UMAP function requires neighborhoods to be computed before computing the UMAP; the UMAP node requires the keys, *neighbors* and *connectivities* to be available to execute correctly. Node effects specify what variables are created by the node and stored in the output data. For example, the UMAP node would store data in the *X_umap* key. Both the node requirements and effects can be affected by modifying node parameters, which makes these unknowable before the user has finalized the pipeline. Instead, we use key strings to identify requirements/effects that are determined from parameters and fetch the relevant values when the user is ready to validate their pipeline.

The method to initialize a node is also used to describe parameters that can be set by the user. This includes the parameter name, default value, and data type. Since the user interacts with the HTML frontend, all parameters are effectively defined as character strings. The parameter type is used to instruct PSCS to convert the character strings to the specified type using the Typomancy^24^ package. Consequently, developers don’t need to code their own input handling. If needed, parameters can be processed before being stored in the node for later use.

Lastly, the node’s *run* method is the core purpose of all nodes and has three steps. First, get the node’s input data from the previous node. Second, process the data and store the results in the Annotated Data object. Third, make the data available to following nodes. The first and third steps are standard and rarely need modification. When the whole pipeline is executed, the input nodes are identified and load their data from disk. From there, nodes are executed in sequence, with the Annotated Data being passed by reference. Passing by reference and storing results in the same object allows us to minimize memory requirements; in cases where a node has multiple outputs, the data is duplicated, and each duplicate is then passed by reference. This is done to avoid possible race conditions, where separate paths might write to the same key in the Annotated Data object.

### Input Data and Uploads

After processing raw mass spectrometry data to quantify peptides and proteins, users can upload the tabular data as .csv files, including sample metadata (e.g., batch number, treatment code) and variable metadata (e.g., whether a column is mitochondrial protein), where it is converted to an Annotated Data^25^ object that is compatible with Scanpy^7,8^. The data is then available for any pipeline, and once published can be imported into other projects.

### Pipeline Execution

Executing the pipeline requires four sets of files: the descriptor file, the data file(s) to be processed, the input mapping file, and the job file for the remote computing resource. The descriptor file specifies the code to use for each node, its parameters, and its connections to other nodes. The data files are specified by the user when the pipeline is being submitted. The input mapping file matches the data files to the pipeline’s input nodes, allowing for pipelines to have multiple inputs. Lastly, the remote computing resources use workload managers (e.g., Slurm^26^, HTCondor^27^, etc.) which need a brief script to schedule computing jobs. The four sets of files are sent to the remote resource, the pipeline is constructed from the descriptor, and the analysis is executed in a Singularity container. Once the analysis is complete, the results are transferred back to the PSCS server, registered into the database, and then made available to the user.

### External App Integration

Interactive apps relevant to single cell can be integrated into PSCS as a microservice. As an example, pipelines can use the CellXGene node in the pipeline designer to export their processed data and make it compatible with CellXGene Annotate^18^. Once available, users can load their data into CellXGene and view a 2D projection (e.g., UMAP^28^, tSNE^29^, principle components), of samples and distributions of observational metadata. New apps can be integrated into PSCS with relative ease, assuming that their resource requirements can be met.

