## Supplementary figures for "Distributed Collaboration for Data, Analysis Pipelines, and Results in Single-Cell Omics"

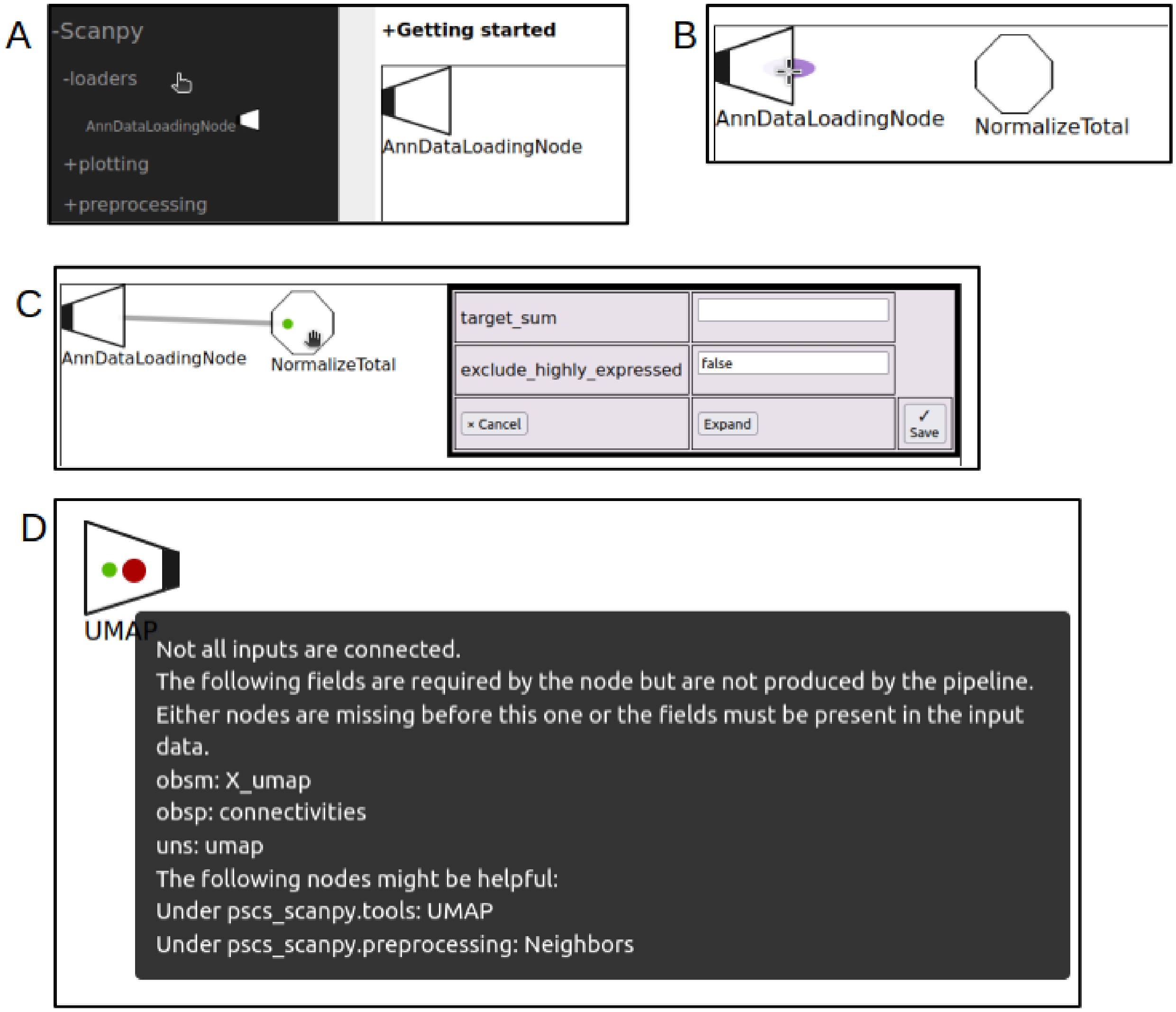


**Supplemental Figure 1. Parts of the pipeline designer. (A)** Analytical packages are used to select the nodes to add to analysis. **(B)** Pipelines are constructed by connecting one node's output port to another node's input by dragging the connection. **(C)** Node parameters can be set by double-clicking a node. A full list of parameters can be viewed under the **Expand** button. **(D)** The pipeline validator identifies problems in the analysis that need to be addressed. If a node's requirements are not met, the validator suggests nodes that could resolve the problem.

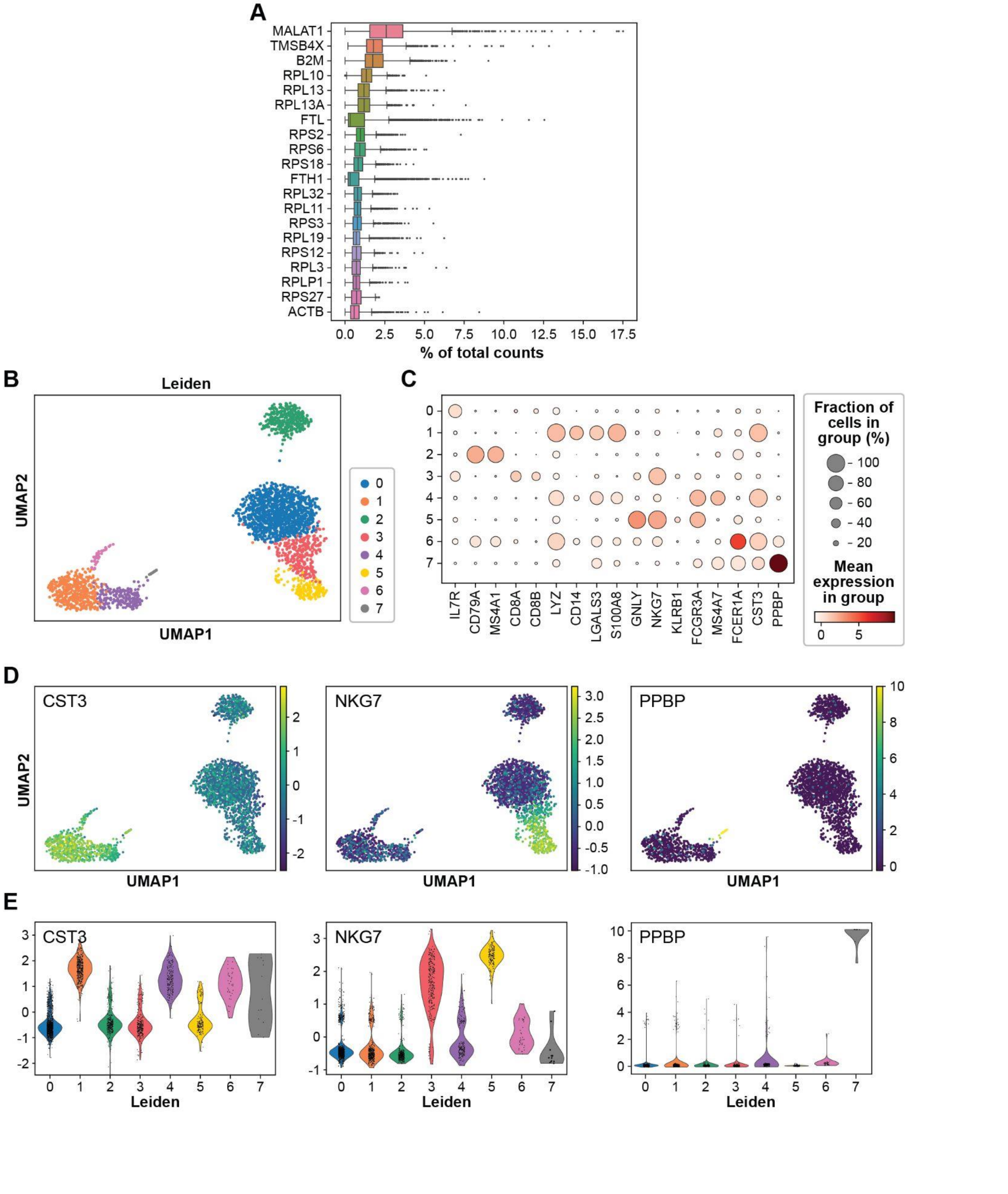


**Supplemental Figure 2. Scanpy PBMC 3k tutorial results using PSCS. (A)** Pre-processing protein distribution. **(B)** UMAP with Leiden clusters. **(C)** Dotplot of the protein distribution across Leiden clusters. **(D)** Protein quantities in cells overlaid on the UMAP. **(E)** Distribution of proteins in cells divided by Leiden clusters.
